# Stress-induced alterations of mesocortical and mesolimbic dopaminergic pathways

**DOI:** 10.1101/2021.01.04.425341

**Authors:** F Quessy, T Bittar, LJ Blanchette, M Lévesque, B Labonté

## Abstract

Our ability to develop the cognitive strategies required to deal with daily-life stress is regulated by region-specific neuronal networks. Experimental evidences suggest that prolonged stress in mice induces depressive-like behaviors via morphological, functional and molecular changes affecting the mesolimbic and mesocortical dopaminergic pathways. Yet, the molecular interactions underlying these changes are still poorly understood and whether they affect males and females similarly is unknown. Here, we used chronic social defeat stress (CSDS) to induce depressive-like behaviors in male and female mice. Density of the mesolimbic and cortical projections was assessed via immuno-histochemistry combined with Sholl analysis along with the staining of the activity-dependent markers pERK and c-Fos in the ventral tegmental area (VTA), nucleus accumbens (NAc) and medial prefrontal cortex (mPFC). We showed that social stress decreases the density of dopaminergic axonal projections to the mPFC but not to the NAc in susceptible and resilient mice. This was accompanied by sex-specific alterations of pERK and c-Fos expression in the VTA of susceptible but not resilient mice. Our results indicate that social defeat stress impacts the mesolimbic and mesocortical pathways by altering the molecular interactions regulating somatic and axonal plasticity differently in males and females.

## Introduction

Major depressive disorder (MDD) is a complex and highly heterogenous mental disorder affecting yearly more than 267 million people worldwide^1^. This heterogeneity, combined with complex environmental, genetic and molecular etiologies, interferes with our capacity to treat MDD efficiently and consequently imposes major economic and medical burden on modern societies^2^. Part of the complexity of MDD emerges from its sexually dimorphic nature. In Canada, the incidence of MDD is 1.7-fold greater in women compared to men^3^. Clinical studies report women to exhibit higher scores of depression, younger age of onset, higher number of depressive episodes and relapse rate^4–7^. Furthermore, differences in treatment responses to antidepressants have been reported between men and women with MDD^8,9^. Together, this suggests that the functional and molecular mechanisms underlying the expression of the disease may differ significantly in men and women although the nature of these differences and their contribution to the expression of MDD in both sexes remain poorly understood.

Historically, the implication of monoamines, including dopamine (DA) has dominated the field of MDD research showing the key roles DA circuits on mood and motivation^10^. DA neurons in the brain are mainly found in the ventral tegmental area (VTA) and the substantia nigra pars compacta (SNpc). There is a large body of evidence showing molecular, functional and transcriptional alterations of the DA system induced by chronic stress. For instance, human studies associated single nucleotide polymorphisms (SNPs) on dopaminergic genes with elevated risk to develop MDD^11,12^. Lower expression of the dopamine transporter (DAT) has also been reported in the brain of MDD patients^13,14^ and mice susceptible to social stress^15–17^. Furthermore, impaired binding of DA to dopamine receptor 1 and 2 (DRD) has been reported in the striatum of depressed suicide completers^18^ although no change in DA levels have been reported in suicide completers^19–21^. Importantly, the role DRD2 in mediating susceptibility to social stress has also been confirmed in rodents^22–24^.

Importantly, DA neurons from the VTA project to several parts of the brain to modulate a wide spectrum of emotionally-relevant behavioral processes such as reward, salience, fear, aversion and memory^25–27^ many of which have been shown to be impaired in MDD patients and mouse models of chronic stress^28–30^. Amongst these different pathways, DA projections from the VTA to the nucleus accumbens (NAc) and the medial prefrontal cortex (mPFC) form the mesolimbic and mesocortical DA pathways, respectively. These two pathways have been consistently associated with the expression of stress responses in mice^31–33^. For instance, previous functional studies showed that optogenetic activation of the mesolimbic pathway induces stress susceptibility while its inhibition promotes resilience to social stress^34^. In contrast, inhibition of the mesocortical pathway was shown to induce susceptibility to social stress with no behavioral effect associated with the inhibition of this pathway in male mice^32,35^. Moreover, a study revealed opposite dopamine metabolism responses between mesocortical and mesolimbic pathways in mice undergoing forced swim test^36^. While these results support the distinct contribution of both pathways in mediating the impact of social stress, their contribution in male and female remains unclear.

In this study, we combined morphological and molecular approaches to investigate the effects of chronic social stress on the mesolimbic and mesocortical DA circuits and compared these effects in males and females. Overall, through these approaches, we provide insights into the distinct contribution of both pathways in mediating susceptibility or resilience to social stress in males and females.

## Results

In this study, we compared the impact of chronic social defeat stress (CSDS) on the morphological and molecular properties of the mesolimbic and mesocortical dopaminergic pathways in male and female mice (**Figure 1A**). 10 days of CSDS induced social avoidance in 26 males (70,27%) and 11 females (55%) while the remaining 11 males and 9 females continued to interact with the CD1 target (Male: F_(2,47)_=45.35; *p*<0.0001, Female: F_(2,25)_=16.27; *p*<0.0001, **Figure 1B**). As expected, susceptible male and female mice spent more time in the corners of the open field, avoiding CD1 targets compared to resilient males (F_(2,45)_=4.204; *p*<0.05 and females (F_(2,25)_=8.474; *p*<0.005, **Figure 1C**).

**Figure 1.**
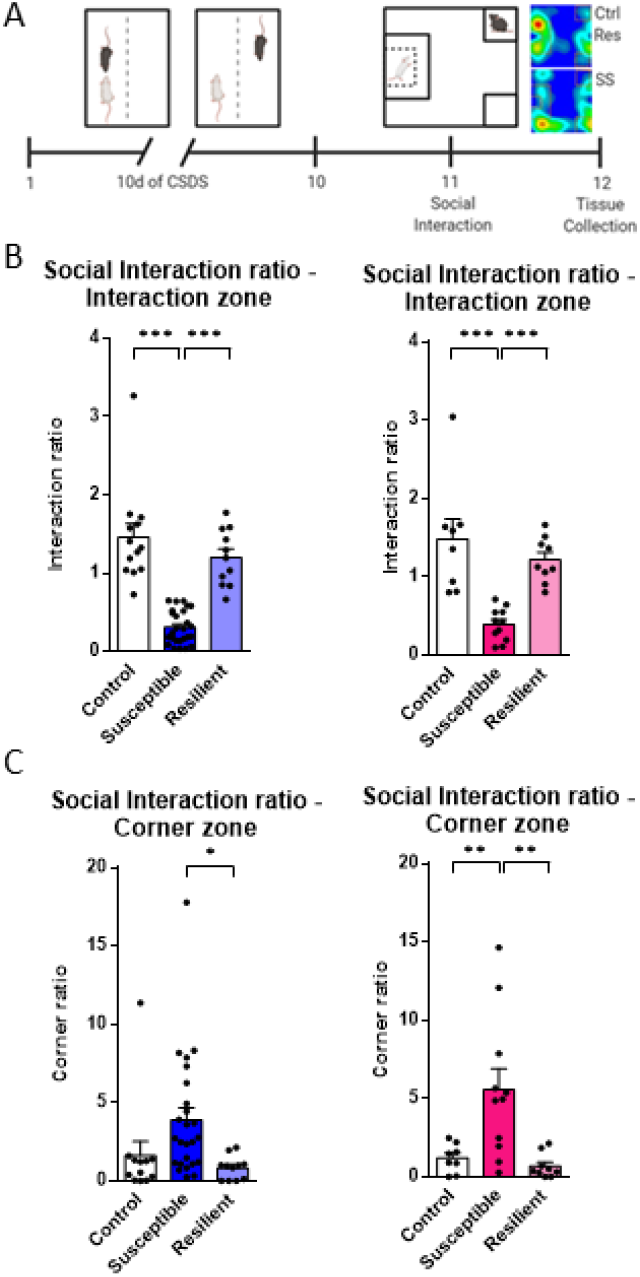
CSDS induces depressive-like social avoidance in males and females. (**A**) Schematic diagram showing the experimental procedure for CSDS. (**B**) Repeated CSDS induces social avoidance in susceptible male (in blue) and female mice (in pink) but resilient mice keep interacting with CD1 aggressors [One-way ANOVA, Male: F_(2,47)_=45.35; *p*<0.0001, Female: F_(2,25)_=16.27; *p*<0.0001), Control (M/F): n=13/8, Susceptible (M/F): n=26/11, Resilient (M/F): n=11/9]. (**C**) Corner ratio results for male and female mice [One-way ANOVA, Male: F_(2,45)_=4.204; *p*<0.05; Female: F_(2,25)_=8.474; *p*<0.005, Control (M/F): n=13/8, Susceptible (M/F): n=26/11, Resilient (M/F): n=11/9]. Bar graphs show mean ± SEM.

### Sex-specific impact of chronic social stress on the morphological properties of DA mesolimbic and mesocortical circuits

Our first objective was to test whether chronic social stress induces a morphological reorganization of DA circuits in males and females. We first tested the impact of CSDS on DA axonal arborization in the mPFC of males and females combining IHC staining with 2D Sholl analysis^37^ to quantify variations in the axonal arborization of VTA DA neurons projecting to the different layers of the mPFC. Consistent with previous studies^38^, our analysis of the mesocortical pathway revealed sparce DA inputs located mostly to the layer V and VI of the mPFC, with axonal ramifications to layers II/III and I (**Figure 2A**). Quantification of cortical DA input shows that susceptible, but not resilient mice, have smaller DA axonal density in the mPFC compared to controls (**SI Table 1 and 2, Figure 2B**). While consistently observed across all cortical layers, these effects are more prominent in layers V/VI of the mPFC (**SI Table 1 and 2**). Additionally, regression analysis revealed a significant positive correlation between social interaction ratios and TH+ axonal arborization in the mPFC (r^2^=0.1213; *p*<0.05**, Figure 2C**) suggesting that smaller arborization of DA inputs in the mPFC may increase susceptibility to CSDS in male and female mice.

**Figure 2.**
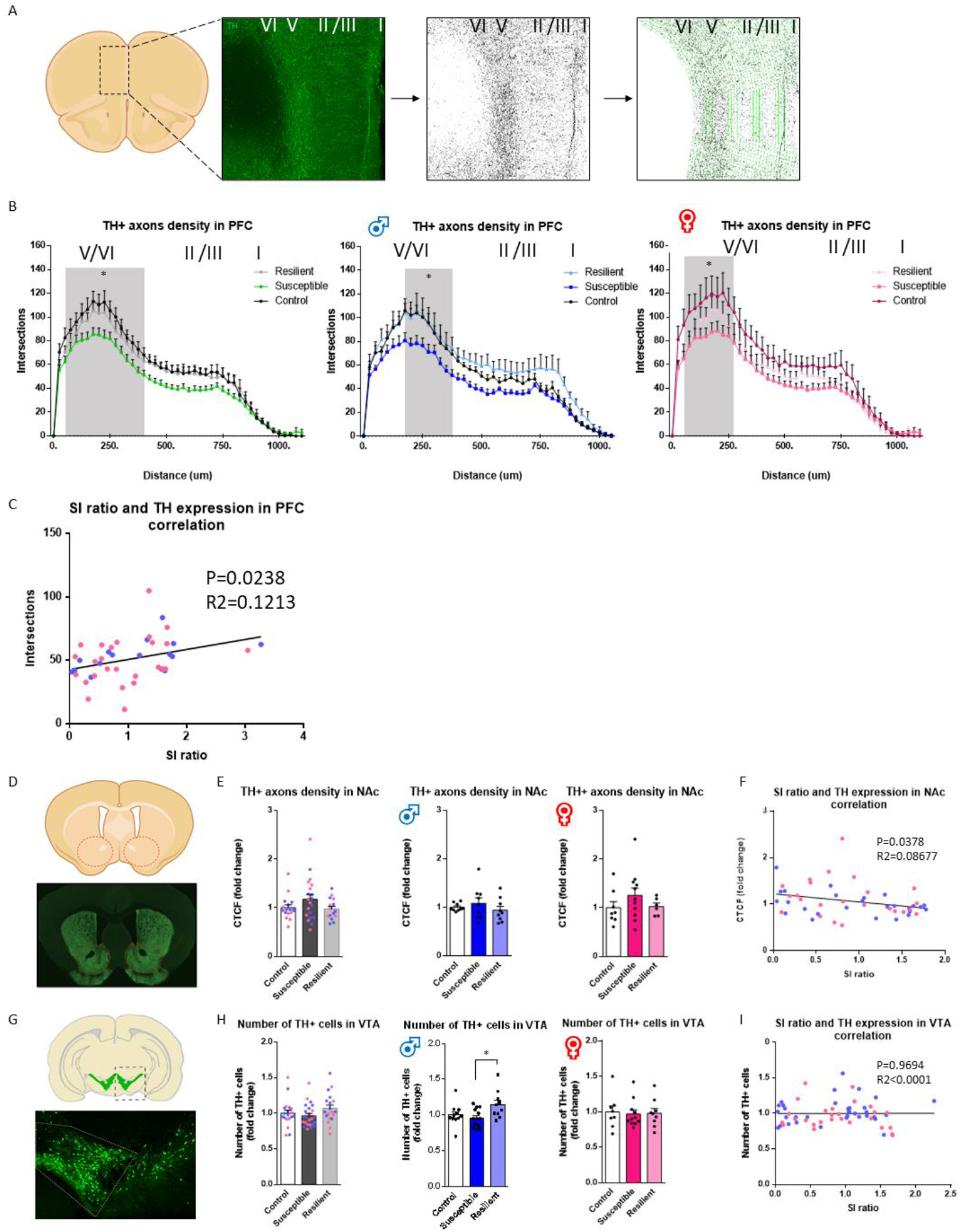
CSDS induces pathway-specific morphological changes in both mesolimbic and mesocortical circuits. (**A**) Schematic diagram showing TH+ staining in the mPFC. (**B**) TH+ neurite density repartition in the different layers of the mPFC. Distance 0 correspond to the corpus collosum (Layer VI) up to Layer I [Two-way ANOVA with Tukey’s multiple comparisons test significant between control and susceptible between 50um to 400um combining males and females (left panel), 175um to 325um for males (middle panel) and 50um to 275um for females. Control (M/F): n=16/7, Susceptible (M/F): n=5/12, Resilient (M/F): n=5/6]. (**C**) Correlation between SI ratios and TH expression in the mPFC [linear regression: R^2^=0.121, *p*<0.05; n=41]. (**D**) Schematic diagram showing TH+ staining in the NAc. (**E**) Fluorescence density of TH+ axons in the NAc (females in pink and males in blue [One-way ANOVA, Males and females combined: F_(2,49)_=2.038; *p*=n.s., Males: F_(2,24)_=0.7369; *p*=n.s., Female: F_(2,22)_=1.098; *p*=n.s., Control (M/F): n=9/8, Susceptible (M/F): n=9/11, Resilient (M/F): n=9/6]). (**F**) Correlation between SI ratio and CTCF values in NAc [linear regression: r^2^=0.087, *p*<0.05; n=52]. (**G**) Schematic diagram showing TH staining in the VTA. (**H**) Stereological counts of TH+ neurons in VTA (females in pink and males in blue [One-way ANOVA, Males and females combined (left panel): F_(2,59)_=1.656; *p*=n.s., Males (middle panel): F_(2,31)_=4.129; *p*<0.05, Females (right panel): F_(2,25)_=0.04272; *p*=n.s, Control (M/F): n=11/8, Susceptible (M/F): n=13/11, Resilient (M/F): n=10/9]). (**I**) Correlation between SI ratio and the number of TH+ cells in the VTA [linear regression: R^2^<0.0001, *p*=n.s.; n=61]. Bar graphs show mean ± SEM. Data are represented as fold change from same group control values. Each dot represents one mouse. Values derived from at least three sections per brain. Scale bar, 100um.

We next measured DA innervation in the NAc. Because DA inputs to the NAc are too dense to perform Sholl analysis, we quantified DA inputs by measuring corrected total cell fluorescence intensity (CTCF^39–41^; **Figure 2D**). Our analysis indicates that dopaminergic axonal density in the entire NAc is not affected by chronic social stress in either males or females (**Figure 2E**). However, linear regression analysis revealed a significant negative correlation between social interaction ratios and levels of TH fluorescence in the NAc (r^2^=0.08677; *p*<0.05, **Figure 2F**) suggesting that smaller DA axonal arborization in the NAc may promote resilience to social stress. We then analyzed TH intensity more specifically in the core and shell sub-regions of the NAc since both sub-regions have been shown to be involved in the control of distinct behavioral features relevant to emotional responses to chronic stress^42–44^. In the shell, we found a trend toward a higher TH fluorescence levels in mice susceptible to CSDS compared to controls when combining males and females (F_(2,49)_=2.789; *p*<0.1, **SI Figure 1A**). We found no significant difference in the core region (**SI Figure 1C**). Linear regression revealed a significant negative correlation between social interaction ratios and TH density in the shell (r^2^=0.08857; *p*<0.05, **SI Figure 1B**) with no correlation in the core subregion of the NAc (**SI Figure 1D**). Finally, no change in TH intensity was observed in mice resilient to CSDS in either NAc shell or core sub-regions. Together, our results suggest that increased DA axonal arborization in NAc shell may increases susceptibility to chronic social stress in mice.

We next counted TH-stained neurons in the VTA (**Figure 2G**) to verify whether changes in DA axonal innervation described above could be due to a change in the number of DA neuron in the VTA. Our analysis indicates that the number of TH+ neurons in the VTA remains unchanged in either susceptible and resilient compared to control mice (**Figure 2H**) with no association between social interaction ratios and the number of TH+ neurons in the VTA (**Figure 2I**). In sum, our analysis suggests that CSDS induces opposite reorganization of the morphological properties of mesolimbic and mesocortical DA pathways, with larger density of DA inputs to the NAc, and lower DA arborization in the mPFC promoting susceptibility to social stress in male and female mice.

### Social stress affects the ERK pathway of susceptible mice

Our next objective was to test whether the morphological remodeling of the mesolimbic and mesocortical DA pathways induced by CSDS associates with functional changes in the activity of target brain regions by quantifying the expression of the intracellular activity markers phospho-ERK (pERK) and c-Fos in the VTA, the NAc and mPFC of susceptible and resilient male and female mice.

We first quantified the number of pERK+ and c-fos+ neurons within TH+ DA neurons of the VTA (**Figure 3A**). Our sex-specific analyses revealed a significant downregulation of pERK expression in TH+ DA neurons from susceptible male mice compared to controls (F_(2,19)_=5.071; P<0.05, **Figure 3B**) with no change in susceptible females (**Figure 3B**). We found no change in both males and females from the resilient groups compared to controls. Regression analysis also failed to reveal any significant correlation between social interaction ratios and pERK expression in TH+ dopaminergic neurons (**Figure 3C**). Similarly, we found no change in the expression of c-Fos in TH+ DA neurons from the VTA (**Figure 4A**) although a trend toward a significant increase was found in females susceptible to CSDS compared to controls (F_(2,21)_=3.384; *p*=0.053, **Figure 4A**).

**Figure 3.**
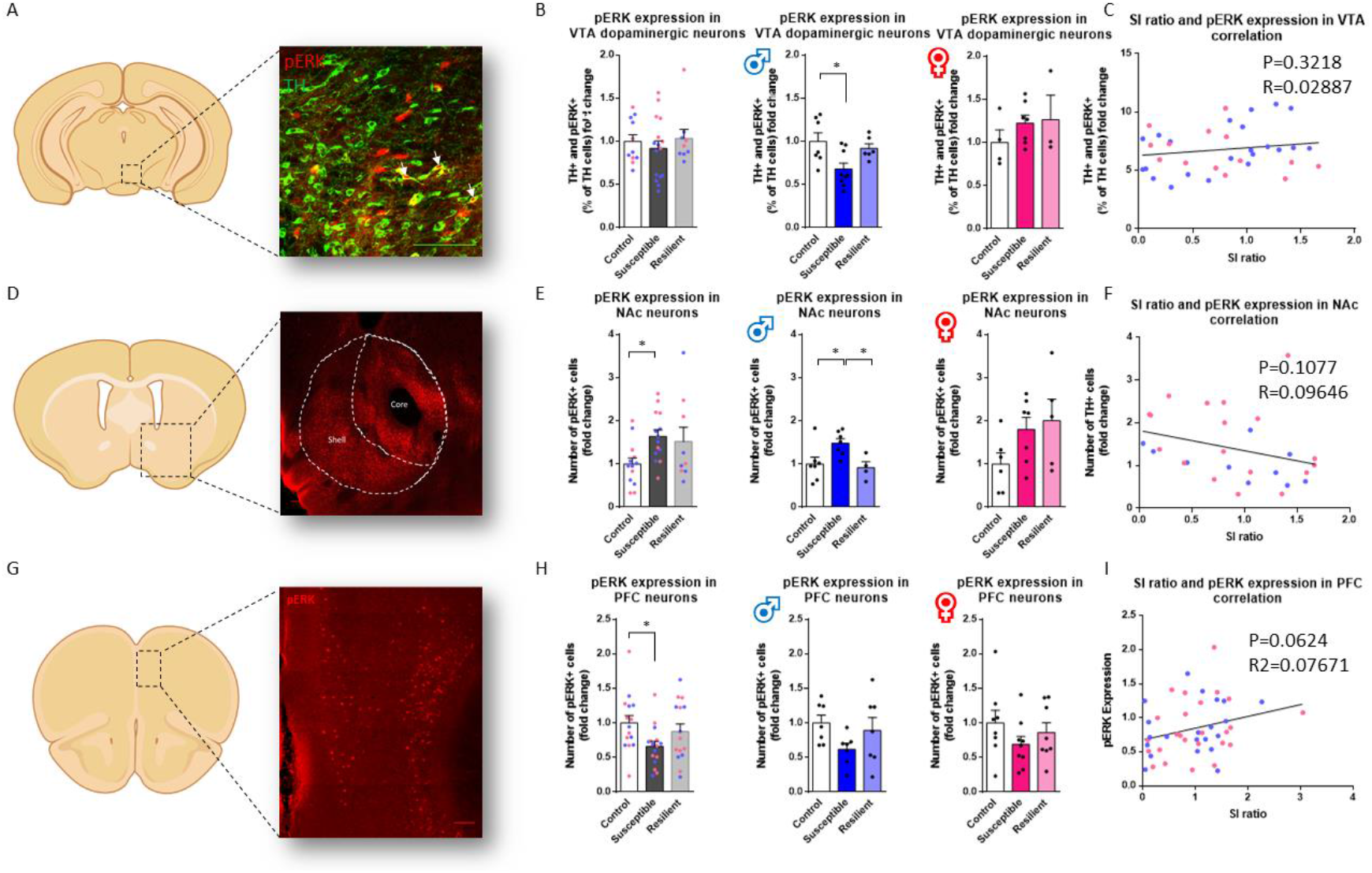
CSDS induces region-specific changes in pERK expression in males and females. (**A**) Schematic diagram showing pERK (red) and TH (green) staining in the VTA. (**B**) Stereological count of TH+ and pERK+ neurons in the VTA (females in pink and males in blue [One-way ANOVA, males and females combined: F_(2,33)_=0.4463; *p*=n.s, Males: F_(2,19)_=5.071; *p*<0.05, Females: F_(2,11)_=0.8122; *p*=n.s, Control (M/F): n=7/4, Susceptible (M/F): n=9/7, Resilient (M/F): n=6/3]). (**C**) Correlation between SI ratios and the number of TH+/pERK+ cells in the VTA [linear regression: R^2^=0.029; *p*=n.s, n=36]. (**D**) Schematic diagram showing pERK+ neurons in the NAc. (**E**) Stereological count of pERK+ neurons in NAc (females in pink and males in blue [One-way ANOVA, Males and females combined (left panel): F_(2,33)_=3.254; *p*<0.1, Males (middle panel): F_(2,15)_=4.932; *p*<0.05, Females (right panel): F_(2,15)_=2.317; *p*=n.s, Control (M/F): n=7/6, Susceptible (M/F): n=7/7, Resilient (M/F): n=4/5]). (**F**) Correlation between SI ratios and the number of pERK+ cells in the NAc [linear regression: r^2^=0.096, *p*=n.s; n=28]. (**G**) Schematic diagram showing pERK staining (red) in the mPFC. (**H**) Stereological count of pERK+ neurons in mPFC (females in pink and males in blue [One-way ANOVA, Males and females combined (left panel): F_(2,43)_=3.089; *p*<0.1, Males (Middle panel): F_(2,18)_=2.075; *p*=n.s, Females (Right panel): F_(2,22)_=1.103; *p*=n.s, Control (M/F): n=7/8, Susceptible (M/F): n=7/9, Resilient (M/F): n=7/8]). (**I**) Correlation between SI ratios and the number of pERK+ cells in the NAc [linear regression: r^2^=0.077, *p*<0.1; n=46]. Bar graphs show mean ± SEM. Data are represented as fold change from same group control values. Each dot represents one mouse. Values derived from at least three sections per brain. Scale bar, 100um.

**Figure 4.**
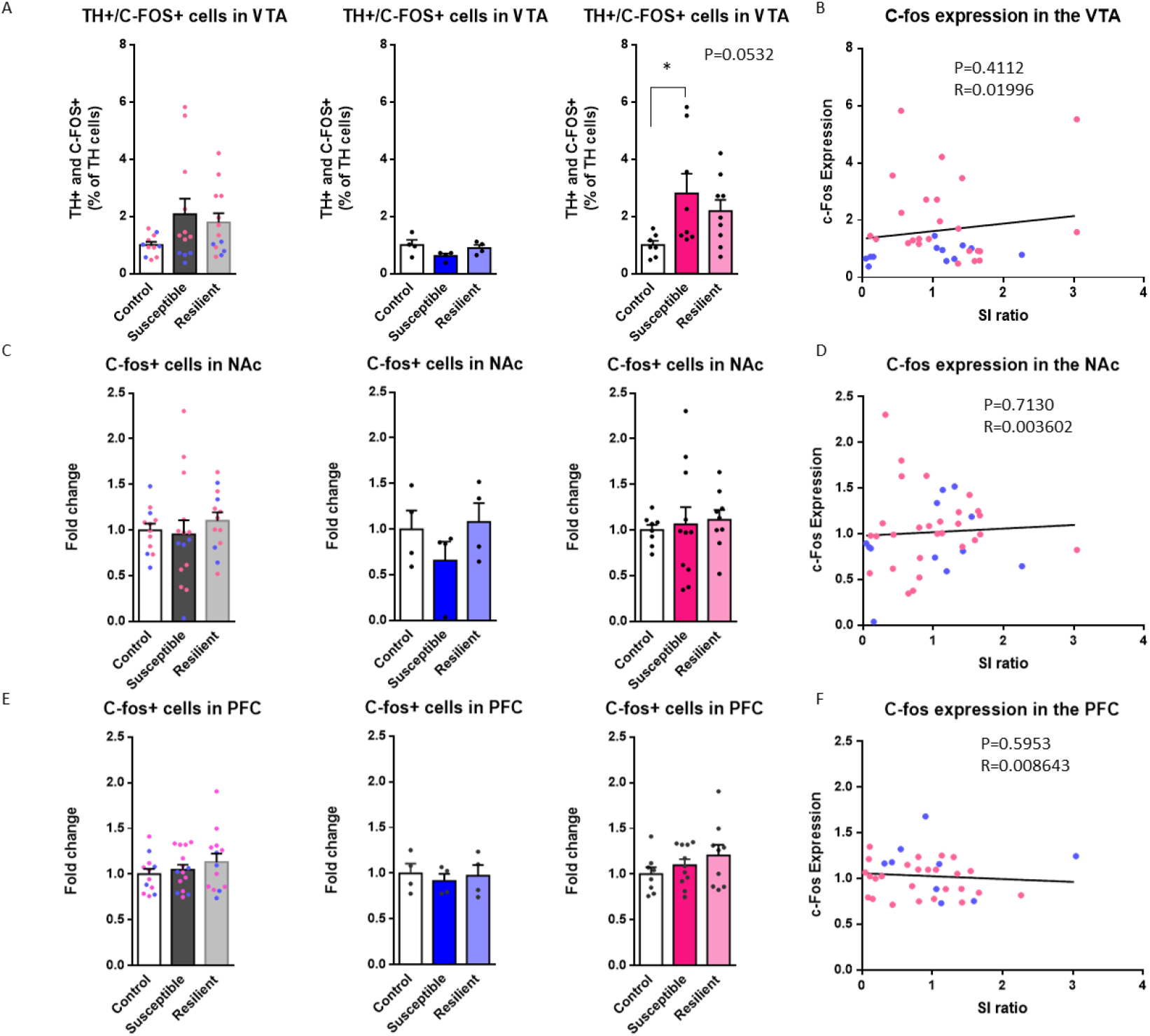
CSDS induces region-specific changes in c-Fos expression in males and females. (**A**) Stereological counts of TH+/c-Fos+ neurons in the VTA (females in pink and males in blue [One-way ANOVA, Males and females combined (left panel): F_(2,33)_=2.069; *p*=n.s., Males (middle panel): F_(2,9)_=2.245; *p*=n.s., Females (right panel): F_(2,21)_=3.384; *p*<0.1, Control (M/F): n=4/7, Susceptible (M/F): n=4/8, Resilient (M/F): n=4/9]. (**B**) Correlation between SI ratios and the number of TH/c-Fos+ cells in the VTA [linear regression: r^2^=0.020, *p*=n.s., n=36]. (**C**) Stereological counts of c-Fos+ neurons in the NAc (females in pink and males in blue [One-way ANOVA, Males and females combined (left panel): F_(2,37)_=0.4174; *p*=n.s., Males (middle panel): F_(2,9)_=1.176; *p*=n.s, Females (right panel): F_(2,25)_=0.1415; *p*=n.s., Control (M/F): n=4/8, Susceptible (M/F): n=4/11, Resilient (M/F): n=4/9]). (**D**) Correlation between SI ratios and the number of c-Fos+ cells in the NAc [linear regression: R2=0.004; *p*=n.s., n=40] (**E**) Stereological counts of c-fos+ neurons in the mPFC (females in pink and males in blue [One-way ANOVA, Males and females combined (left panel): F_(2,37)_=0.8702; *p*=n.s., Males (middle panel): F_(2,9)_=0.1813; *p*=n.s., Females (right panel): F_(2,25)_=1.219; *p*=n.s., Control (M/F): n=4/8, Susceptible (M/F): n=4/11, Resilient (M/F): n=4/9]). (**f**) Correlation between SI ratios and the number of c-Fos+ cells in the mPFC [linear regression: r^2^=0.009; *p*=n.s, n=35]. Bar graphs show mean ± SEM. Data are represented as fold change from same group control values. Each dot represents one mouse. Values derived from at least three sections per brain. Scale bar, 100um.

In the NAc, our analysis revealed a significant increase in pERK in male (F_(2,15)_=4.932; *p*<0.05) but not female susceptible mice(**Figure 3D-E**). Interestingly, our results suggest that increased pERK expression is common to the NAc shell and core, both regions showing significant upregulation of pERK expression in susceptible mice compared to controls (Shell: F_(2,33)_=3.470; *p*<0.05; Core total: F_(2,33)_=3.719; *p*<0.05) although sex-specific effect were found in males and females (Shell male: n.s.; Core male: F_(2,20)_=5.340; *p*<0.05; Shell female: F_(2,15)_=3.779; *p*<0.05; Core female: n.s., **SI Figure 2**). Our regression analysis revealed no significant correlation between social interaction ratios and pERK expression in the NAc (**Figure 3F**). In contrast, we found no change in the expression of c-Fos in either susceptible or resilient male and female mice compared to controls (**Figure 4B**).

Finally, analysis of the mPFC revealed a trend toward a significant downregulation of pERK expression in susceptible compared to control mice (F_(2,43)_=3.089; P=0.056) with no effects in males and females specifically (**Figure 3H**). Regression analysis identified a trend toward a significant positive correlation between social interaction ratios and pERK expression suggesting that low levels of pERK expression in the mPFC may promote susceptibility to social stress in males and females (r^2^=0.077; *p*=0.062, **Figure 3I**). In contrast, our analysis revealed no change in the expression of c-Fos in either susceptible or resilient male and female mice compared to controls, **Figure 4C**). Interestingly, further analyses revealed that the downregulation of pERK in susceptible mice is specific to pyramidal neurons from superficial layers of the infralimbic mPFC (Superficial layers: F_(2,30)_=3.814; *p*<0.05; infralimbic: F_(2,43)_=3.258; *p*<0.05) with no effect in the prelimbic mPFC (**SI Figure 3**). Together, this suggests that CSDS interferes in a sex-specific fashion with ERK but not c-fos intracellular signaling cascades regulating neuronal activity in the target brain regions of the mesolimbic and mesocortical DA pathways with higher levels of pERK in the NAc and lower levels in the mPFC, respectively, may promote the expression of susceptibility to social stress in males and females.

## Discussion

Dysregulation of dopaminergic transmission has been widely involved in the pathophysiology of depression^45^. Depressive disorders symptoms such as social avoidance, anhedonia, lost of appetite, helplessness and amotivation have been consistently associated with dysfunctions of dopamine signalling and functions^35,46–48^. Now, it is increasingly accepted that males and females respond differently to chronic stress^49,50^ suggesting that the DA system could exhibit sex-specific morphological and molecular adaptations to chronic stress in a sex-specific fashion. Here, we report findings showing morphological and molecular alterations in mesocortical and mesolimbic dopaminergic pathways of both stressed male and female mice and provide new insights into the implication of dopaminergic circuits in mediating stress susceptibility in a sex-specific fashion.

Neuroanatomical and morphological changes in several brain regions have been consistently associated with MDD. For instance, post-mortem studies on MDD patients showed a decrease of neuronal soma size in layer V and VI compared to healthy subjects^51,52^. Smaller volume of dorsal striatal gray matter was also shown to correlate with suicidal ideation in adolescent^53^. Similarly, susceptible mice to CSDS exhibit lower spine density in the mPFC and increased spine density in the NAc and the VTA^54^ associated with higher and lower volume in the VTA and NAc, respectively^55^. Here, we provide results showing that CSDS does not change the number of DA cells in the VTA but rather induces an important remodelling of the DA mesolimbic and mesocortical circuits in both sexes. In the NAc, our findings suggest that chronic social stress increases DA axons density in the shell, but not in the core. DA projections form the VTA target mainly the shell part of the NAc composed of MSN expressing either DRD1, DRD2 or both receptors^56,57^. Importantly, lower frequency of excitatory input onto DRD1 and higher frequency onto DRD2 expressing MSNs has been previously reported in the NAc of susceptible mice to CSDS. Accordingly, optogenetic stimulation of DRD1 MSNs was associated with the expression of resilience while the optogenetic inhibition of DRD2 MSNs was shown to induce depressive-like behaviors following CSDS^58^. Accordingly, an increase in projections density in the NAc shell as supported by our results could cause an imbalance in dopamine regulation in this region and drive the behavioral effects of CSDS. More work will be required to determine whether these changes impact DA transmission at DRD1 or DRD2 MSNs in males and females.

In contrast, we found a significant decrease of axonal dopaminergic density in the layer V and VI of mPFC in susceptible mice of both males and females. DA neurons from the VTA project mainly on layer V and VI of the mPFC^59^. Interestingly, optogenetic inhibition of the mesocortical DA pathway was previously shown to promote susceptibility to CSDS in males^35^. Functionally, speaking, the lower density of DA axons observed in our study is likely to decrease DA signalling in this region and promote social withdrawal in males and females after CSDS. Importantly, while providing interesting insights into the morphological impact of CSDS on DA circuits, approach using immunohistochemistry for TH carries limitations that could interfere with the interpretation of our results. For instance, it has been estimated that 10% of TH+ axons in the mPFC originates from axons from the locus coeruleus (LC)^60^. However, mPFC inputs from the LC are mainly located in layers II/III^38^ and as such, may not interfere with our main findings in layers V/VI. Nevertheless, the results of our study suggest that CSDS induces morphological alterations in DA axons and could lead to distinct functional impairments in dopaminergic signalling in the NAc and mPFC.

Importantly, our findings show that morphological changes in DA circuits induced by chronic social stress associate with molecular changes affecting intracellular signalling in DA neurons and their targeted brain regions. Transcriptional profiling studies showed before that chronic stress including CSDS induces a global reorganization of transcriptional structures of DA neurons and its targeted brain regions^61,62^. Importantly, these alterations have also been associated with changes within several intracellular signalling pathways including MAPK^43,63,64^, AKT^65,66^, mTOR^67^ and GTPase pathways^68,69^. Of particular interest, molecular markers such as pERK and c-Fos have been used as proxies of neuronal activity in several mouse models of chronic and acute stress^49,70,71^. Here, we provide evidence for a downregulation of pERK signalling in TH+ neurons of susceptible male mice specifically. This is in contrast with results from previous studies showing elevated ERK activity in neurons of the VTA after CSDS^64,71^. Noticeably, these effects were observed in a small proportion of DA neurons in the VTA (approx. 6%) while most of the pERK+ neurons in the VTA did not express TH which could explain the discrepancies between our study and others. Thus, it is likely that the impact of CSDS on ERK signalling in the VTA may be relevant to the activity of different neuronal populations with distinct neuronal connectivity patterns. In contrast, we found a trend toward an increase in c-Fos expression in the VTA of susceptible female mice which is consistent with previous observations in females after CSDS^49^ and supports the sex-specific involvement of these two intracellular signalling pathways in the processing emotional responses to social cues differently^72^.

Functionally, the pathway-specific morphological changes observed mPFC and NAc projections along with the molecular changes in pERK and c-Fos expression in the VTA following CSDS are likely to impact DA signalling in a pathway specific fashion. Our analysis quantifying pERK and c-Fos expression in the mPFC and NAc support this conclusion although indirectly. In the NAc, our results suggest that CSDS increases pERK levels in the core of male susceptible mice and the shell of susceptible female mice. Previous studies showed that stress activates mainly the shell structure of the NAc^73,74^ while drugs seeking activates the core structure^75^. The pERK expression patterns observed in the core and shell suggest that both structures integrate emotional stressors differently in males and females and suggest that distinct neuronal circuits relevant to these sub-regions in the NAc may contribute differently to the expression of stress in males and females. In the mPFC, our findings support a downregulation of ERK signaling in susceptible mice as shown before after CSDS^76^. It has been shown that ERK activation contribute to upregulation of dendritic spine density in the mPFC and improved learning and memory^76,77^. However, our findings are also in contrast with other studies showing a significant increase in pERK expression in females following chronic variable stress^78^ and suggest that different stress types may impose different impact on the activity of the mPFC in males and females. These discrepancies could be explained in part by the nature of both stress paradigms, CSDS reproducing psychosocial stress while CVS being composed of physical stressors^79^. While these results suggest that the pathways-specific morphological and molecular changes affecting the mesolimbic and mesocortical pathways in males and females may lead to changes in DA signalling in the mPFC and NAc, our results remain indirect and more work will be required to determine the functional impact of these changes in males and females.

To conclude, results from our study provide new insights on the impact of CSDS on dopaminergic sub-circuits and the contribution of mesolimbic and mesocortical pathways on the expression of depressive-like behaviors associating these effects with morphological and molecular alterations in males and females. In doing so, we increase our understanding of the functional and molecular mechanisms from which sexual dimorphism emerges, defining the clinical characteristics of mood disorders. Ultimately, we will be in a better position to develop therapeutic approaches more suited to treat depression in men and women.

## Methods

### Animals

C57bl6 male (*n*= 50) and female (*n*=28) mice aged 7-8 weeks old and CD1 males aged 4-6 months old were obtained from Charles River Laboratories (USA). In total 50 males and 28 females were subjected to social defeat. Mice were housed under a 12 hours light/dark cycle at 22-24°C with no water and food restriction. CD1 mice were individually housed except during social defeats. All other mice were group-housed (4/cage) before and singly housed after social defeats. All experimental procedures were approved by Laval’s University Institutional Animal Care Committee in respect with the Canadian Council on Animal Care guidelines.

### Chronic Social Defeat Stress

CSDS was performed as described before in males^80^ and females^81^. Briefly, in males, C57bl6 were first introduced to unknown aggressive retired CD1 breeders for a period of 5 minutes during which they were attacked by the resident mice. Following this initial phase, C57bl6 mice were moved to the other side of the divider for 24 hours allowing a continuous sensory contact with the CD1 mouse without physical harm. The same procedure was repeated over 10 days with new unknown CD1 mice every day.

A very similar approach was used in females with the difference that the base of the tail and the pubis of the C57bl6 female mice were soaked with 30 μl of male urine before each defeat bouts. All C57bl6 female mice were associated with a different male urine so that CD1 mice never encountered the same urine twice. Urine collection was performed 1-week preceding social defeat stress with metabolic cages (Tecniplast Group, Tecniplast Canada, Canada). Male mice were housed in metabolic cages overnight. Urine was collected each morning and stored at 4°C after being filtered. The 5-minutes social stress bouts were interrupted every time CD1 male mounted the C57bl6 female mice. Both male and female controls were housed 2/cage separated by a Plexiglas divider and housed in the same room.

### Social interaction

Social-avoidance behavior was assessed with the social interaction test (SI) 24 hours after the end of the social defeat paradigm as described before^80,81^. One hour before the social interaction test, C57bl6 male and female mice were habituated to the testing room illuminated by red light. Briefly, SI test consisted of two phases of 150 seconds each. In the first phase, C57bl6 mice explored the arena with no target CD1 aggressor in the social interaction zone. This initial phase was followed by a second exploratory phase but this time in the presence of an unknown target CD1 aggressor maintained into a mesh-wired enclosure within the social interaction zone. Time spent in the different zones of the arena was automatically recorded through ANY-Maze 4.99 using a top-view camera (ANY-Maze Video Tracking Software, Stoelting Co., USA). Based on social interaction ratios (time in interaction zone with social target/time in interaction zone without social target), defeated mice were designated as susceptible or resilient: susceptible ratio < 1.0; resilient ratio > 1.0. This measure of susceptibility versus resilience has been shown to correlate with other defeat-induced behavioral abnormalities such as anhedonia (e.g. decrease in sucrose preference) and an increased sensitivity to inescapable stress^82–84^. Following the social interaction test, C57bl6 mice were housed individually for 24 hours before sacrifice and tissue collection.

### Perfusions

For morphological and molecular analyses, mice were anesthetized with a lethal dose of 20% (w/v) urethane (Sigma, 51-79-6) and perfused with 40 ml of 0.1 M/L phosphate-buffered saline and 40 ml of 4% (w/v) PFA (Sigma, 30525-89-4) 24h after social interaction. Brains were post-fixed for 24h in PFA 4% and cryoprotected in sucrose 30% before being stored in OCT (Tissue-Tek®, 4583) at −80°C. Brains were sliced on a Leica CM1900 cryostat at 40 μm.

### Immunohistochemistry

Free-floating sections were washed with PBS and then blocked with 1% normal donkey serum and 0.2% Triton X-100 for 2 hours. Brains sections were incubated overnight at 4°C with polyclonal antibodies against tyrosine hydroxylase (TH) 1:500 (Pel-Freez Biologicals, P60101), Phospho-p44/42 MAPK 1:250 (Erk1/2) (Thr202/Tyr204) (Cell Signaling, 9101), DAT 1:500 (Millipore, MAB369) or C-FOS 1:500 (Abcam, Ab190289). The next day, after washes, brain sections were incubated in corresponding secondary antibodies (Alexa Fluor 488 Donkey anti-rabbit IgG 1:400, Life Technologies A-21206; Alexa Fluor 555 Donkey anti-sheep IgG 1:400, Life Technologies A-21099; Alexa Fluor 647 Donkey anti-rabbit IgG 1:400, Life Technologies A-31573) for 1 hour at room temperature. Sections were mounted with anti-fade solution, including 40,6-diamidino-2-phenylindole (Abcam, Ab104139). A Zeiss LSM700 confocal microscope was used for TH morphological and pERK/c-Fos colocalization expression for VTA analysis, and a Leica DMRB slide scanner was used to acquire the immunofluorescence images for TH expression in NAc.

All immunofluorescence images were analyzed using ImageJ Software^85^. TH, pERK and c-Fos expression was assessed via counts for each region of interest (VTA, NAc, mPFC), with a minimum of 3 brain section per mouse. Brain sections were selected using the Allan Brain Atlas^86^. mPFC sections were between 1.42 mm and 2.245 mm of Bregma, NAc between 0.845 mm and 1.545mm of Bregma and VTA between −2.78 mm and −3.38 mm of Bregma. The regions and sub-regions of interest were delimited using Allan Brain atlas reference^86^. TH+, pERK+ and c-Fos+ cells were counted manually using Image J software.^85^ TH density in the NAc and c-Fos expression in the NAc and mPFC was analyzed using corrected total cell fluorescence method (CTCF)^39,41,87^. Expression of TH morphological arborization in prefrontal cortex was assessed using the software Neurite-J 1.1^37^.

### Statistical analysis

Statistical analyses were performed using GraphPad Prism 6.0 (GraphPadSoftware, La Jolla, CA, USA) software. In total, we used 50 male and 28 female mice for analyses. Each point represents one mouse. All data are represented as means ± SEM, and significance is defined as \**p* ≤ 0.05, \*\**p* ≤ 0.01, \*\*\**p* ≤ 0.001.

For social interaction and corner ratios (**Figure 1**), 50 male mice and 28 females were analyzed. Using Grubb test with criteria of more than 1.96 standard deviations from the mean, one male control mouse was discarded. Normality was accessed using D’Agostino and Pearson omnibus test. One-way ANOVA was performed with Tukey’s multiple comparison post hoc tests. Each point represents one mouse. All data are represented as mean ± SEM and significance is defined as \**p*≤0.05, \*\**p*≤0.01, \*\*\**p*≤0.001.

For TH density analysis in the mPFC (**Figure 2A-B**), 16 male mice and 26 females were analyzed. Using Grubb test on the mean of all distances with criteria of more than 1.96 standard deviations from the mean, one control female was discarded. Normality of data was accessed using D’Agostino and Pearson omnibus test. Two-way ANOVA with Tukey’s multiple comparison post hoc test was used to compare group differences. All data are represented as mean ± SEM, and significance is defined as \**p*≤ .05, \*\**p*≤0.01, \*\*\**p*≤0.001. For the regression between the number of intersection and the SI ratio, the mean of intersections was compared to the SI ratio of the respective mice (**Figure 2C**). A linear regression between these two values was performed. Each point represents one mouse with at least three sections of mPFC.

For TH density analysis in the NAc (**Figure 2D-E**), 27 males and 25 females were analyzed. No outliers were identified. Normality of data was accessed using Shapiro-Wilk test. One-way ANOVA with Tukey’s multiple comparison post hoc test and Kruskal-Wallis statistic test with Dunn’s multiple comparison were performed. Each point represents one mouse with at least three sections of the regions. The mean of all sections was used to generate each point. The data were generated from 5 different groups of social defeat. All data are expressed in fold change over respective control groups. All data are represented as mean ± SEM and significance are defined as \**p*≤0.05, \*\**p*≤0.01, \*\*\**p*≤0.001. A linear regression was performed between TH density in the NAc and SI ratios (**Figure 2F**). Each point represents one mouse with at least three sections of NAc.

For TH+ cells analysis in the VTA (**Figure 2G-H**), 22 males and 28 females were analyzed. No outliers were identified. Normality of data was accessed using Shapiro-Wilk test. One-way ANOVA with Tukey’s multiple comparison post hoc test and Kruskal-Wallis statistic test with Dunn’s multiple comparison were performed to compare group differences. Each point represents one mouse with at least three sections of the region. The mean of all sections was used to generate this point. The data were generated from four different groups of social defeat. All data are expressed in fold change over their respective control groups. All data are represented as mean ± SEM and significance is defined as \**p*≤0.05, \*\**p*≤0.01, \*\*\**p*≤0.001. A linear regression was performed between TH density in the VTA and SI ratios (**Figure 2F**). Each point represents one mouse with at least three sections of VTA.

For pERK analysis in the VTA (**Figure 3A-B**), 22 males and 14 females were analyzed. No outliers were identified. Normality of data was accessed using Shapiro-Wilk test. One-way ANOVA was performed with Dunn’s multiple comparison post hoc tests. Each point represents one mouse with at least three sections of the region. The mean of all sections was used to generate this point. The data were generated from 4 different groups of social defeat. All data are expressed in fold change over their respective control groups. All data are represented as mean ± SEM and significance is defined as \**p*≤0.05, \*\**p*≤0.01, \*\*\**p*≤0.001. A linear regression was performed between pERK/TH colocalization in the VTA and SI ratios (**Figure 3C**). Each point represents one mouse with at least three sections of VTA.

For pERK expression analysis in the NAc (**Figure 3D-E**), 18 males and 18 females were analyzed. No outliers were identified. Normality of data was accessed using D’Agostino and Pearson omnibus test. One-way ANOVA was performed with Tukey’s multiple comparison post hoc tests. Each point represents one mouse with at least three sections of the region. The mean of all sections was used to generate this point. The data were generated from 4 different groups of social defeat. All data are expressed in fold change over their respective control groups. All data are represented as means ± SEM and significance are defined as \**p*≤0.05, \*\**p*≤0.01, \*\*\**p*≤0.001. A linear regression was performed between pERK expression in the NAc and SI ratios (**Figure 3F**). Each point represents one mouse with at least three sections of NAc.

For pERK expression analysis in the mPFC (**Figure 3G-H**), 21 males and 25 females were analyzed. No outliers were identified. Normality of data was accessed using Shapiro-Wilk test. One-way ANOVA was performed with Tukey’s multiple comparison post hoc tests. Each point represents one mouse with at least three sections of the region. The mean of all sections was used to generate this point. The data were generated from 4 different groups of social defeat. All data are expressed in fold change over their respective control groups. All data are represented as means ± SEM and significance are defined as \**p*≤0.05, \*\**p*≤0.01, \*\*\**p*≤0.001. A linear regression was performed between pERK expression in the PFC and SI ratios (**Figure 3F**). Each point represents one mouse with at least three sections of mPFC.

For c-Fos expression analysis in the VTA (**Figure 4A**), 12 males and 24 females were analyzed. No outliers were identified. Normality was accessed using D’Agostino and Pearson omnibus test. One-way ANOVA was performed with Tukey’s multiple comparison post hoc tests. Male mice were analyzed using Kruskal-Wallis test with Dunn’s multiple comparison tests. Each point represents one mouse with at least three sections of the regions. The mean of all sections was used to generate this point. The data were generated from 3 different groups of socially defeated mice. All data are expressed in fold change over their respective control groups. All data are represented as means ± SEM and significance are defined as \**p*≤0.05, \*\**p*≤0.01, \*\*\**p*≤0.001. A linear regression was performed between c-Fos expression in the VTA and SI ratios (**Figure 4B**). Each point represents one mouse with at least three sections of VTA.

For c-Fos expression analysis in the NAc and PFC (**Figure 4C-E**), 12 males and 28 females were analyzed. Normality was accessed using D’Agostino and Pearson omnibus test. No outliers were identified. One-way ANOVA was performed with Tukey’s multiple comparison post hoc tests. Male mice were analyzed using Kruskal-Wallis test with Dunn’s multiple comparison tests. Each point represents one mouse with at least three sections of the region. The mean of all sections was used to generate this point. The data were generated from 3 different groups of socially defeated mice. All data are expressed in fold change over their respective control groups. All data are represented as means ± SEM and significance are defined as \**p*≤0.05, \*\**p*≤0.01, \*\*\**p*≤0.001. A linear regression was performed between c-Fos expression in the NAc/PFC and SI ratio (**Figure 4D-F**). Each point represents one mouse with at least three sections of NAc or mPFC.

## Acknowledgements

A special thanks to Drs. A. Barbeau, K. Abdallah and Veronique Rioux for technical assistance. B.L. holds a Sentinelle Nord Research Chair, is supported by a NARSAD young investigator award, a CIHR (SVB397205), and Natural Science and Engineering Research Council (NSERC; RGPIN-2019-06496 to BL and RGPIN-2018-06262 to ML) grants and receives FRQS Junior-1 salary support; this work was also made possible by resources supported by the Quebec Network on Suicide, Mood Disorders and Related Disorder). ML is a career awardee of the FRQS in partnership with Parkinson Quebec (34974). This work is also supported by a grant from NSERC (RGPIN-2018-06262) to M.L.

## Author contributions

F.Q., B.L. and M.L. conceived the project, designed the experiments, and wrote the manuscript. F.Q. also generated and analyzed all the data with the help of L.J.B. L.J.B. contributed to the activity and qPCR analyses. T.B. contributed to the behavioural experiments. All authors contributed to the preparation of the manuscript.

## Additional Information

### Data availability

The datasets generated during and/or analyzed during the current study are available from the corresponding author on reasonable request.

