## Supplemental Information for "Stress-induced alterations of mesocortical and mesolimbic dopaminergic pathways"

Figure S1

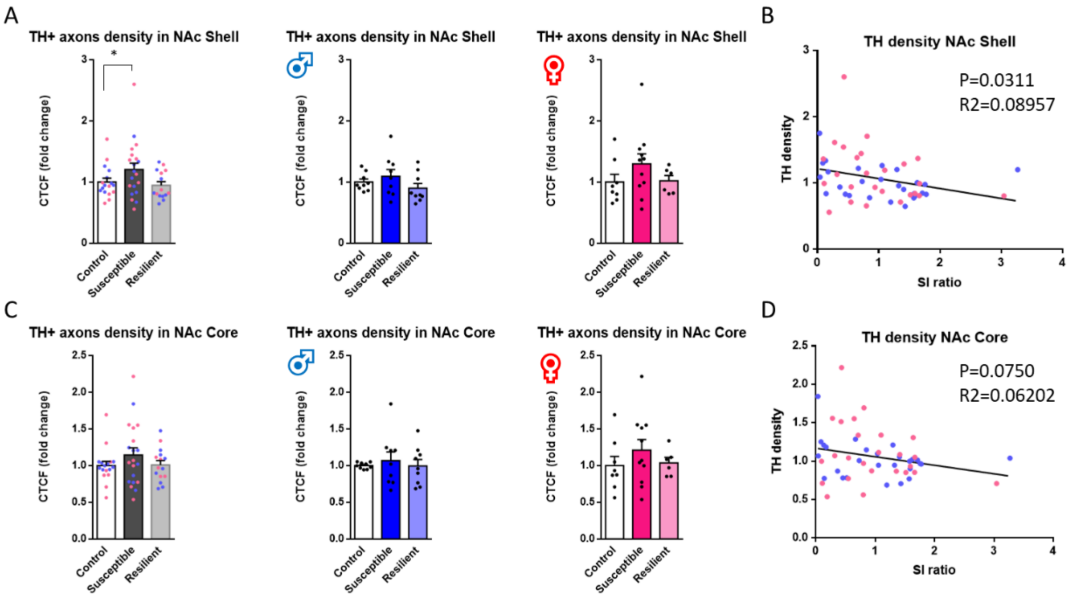


**Figure S1. TH expression in NAc core and shell of males and females after CSDS** (**A**) Fluorescence density of TH+ axons in the NAc shell (females in pink and males in blue [One-way ANOVA: Males and females combined (left panel): $F_{(2,49)}$=2.789; $p$<0.1, Males (middle panel): $F_{(2,24)}$=1.370; $p$=n.s., Females (right panel): $F_{(2,22)}$=1.353; $p$=n.s., Control (M/F): n=9/8, Susceptible (M/F): n=9/11, Resilient (M/F): n=9/6]). (**B**) Correlation between SI ratios and the CTCF values in the NAc shell [linear regression: $R^2$=0.090; p<0.05, n=52]. (**C**) Fluorescence density of TH+ axons in the NAc shell (females in pink and males in blue [One-way ANOVA, males and females combined (left panel): $F_{(2,49)}$=1.174; $p$=n.s., Males (middle panel): $F_{(2,24)}$=0.2215; $p$=n.s., Females (right panel): $F_{(2,22)}$=0.7718; $p$=n.s., Control (M/F): n=9/8, Susceptible (M/F): n=9/11, Resilient (M/F): n=9/6]). (**D**) Correlation between SI ratios and the CTCF values in the NAc core [linear regression: $R^2$=0.062; $p$<0.1, n=52]. Bar graphs show mean ± SEM. Data are represented as fold change from same group control values. Each dot represents one mouse. Values derived from at least three sections per brain.

Figure S2

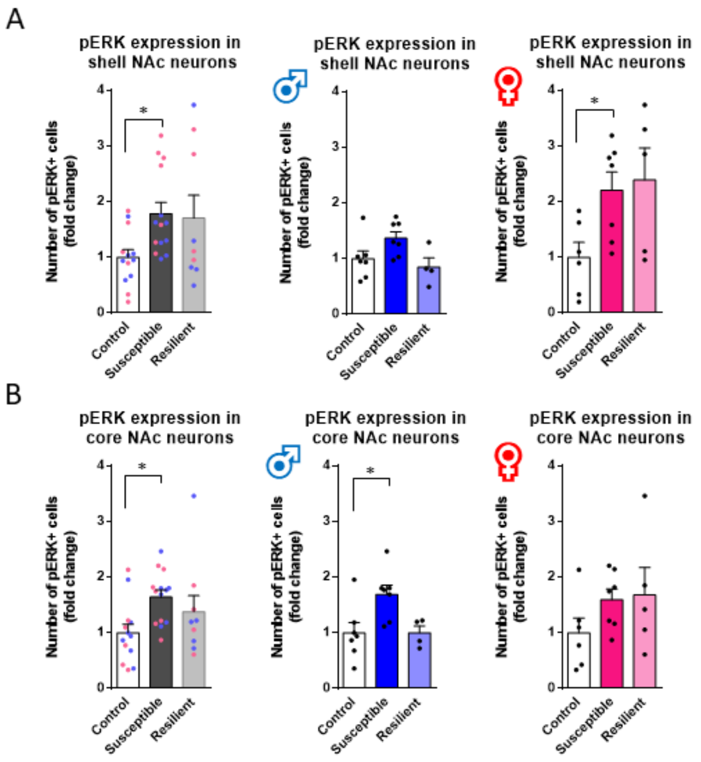


<u>**Figure S2. pERK expression in NAc core and shell of males and females after CSDS**</u> (**A**) Stereological counts of pERK+ neurons in NAc shell (females in pink and males in blue [One-way ANOVA, Males and females combined (left panel): $F_{(2,33)}=3.470$; $p<0.05$, Males (middle panel): $F_{(2,15)}=3.575$; $p<0.1$, Females (Right panel): $F_{(2,15)}=3.779$; $p<0.05$, Control (M/F): n=7/6, Susceptible (M/F): n=7/7, Resilient (M/F): n=4/5]). (**B**) Stereological count of pERK+ neurons in NAc core (females in pink and males in blue [One-way ANOVA, Males and females combined (left panel): $F_{(2,33)}=3.719$; $p<0.05$, Males (middle panel): $F_{(2,24)}=5.340$; $p<0.05$, Females (right panel): $F_{(2,15)}=0.1.422$; $p$=n.s., Control (M/F): n=7/6, Susceptible (M/F): n=7/7, Resilient (M/F): n=4/5]). Bar graphs show mean ± SEM. Data are represented as fold change from same group control values. Each dot represents one mouse. Values derived from at least three sections per brain.

Figure S3

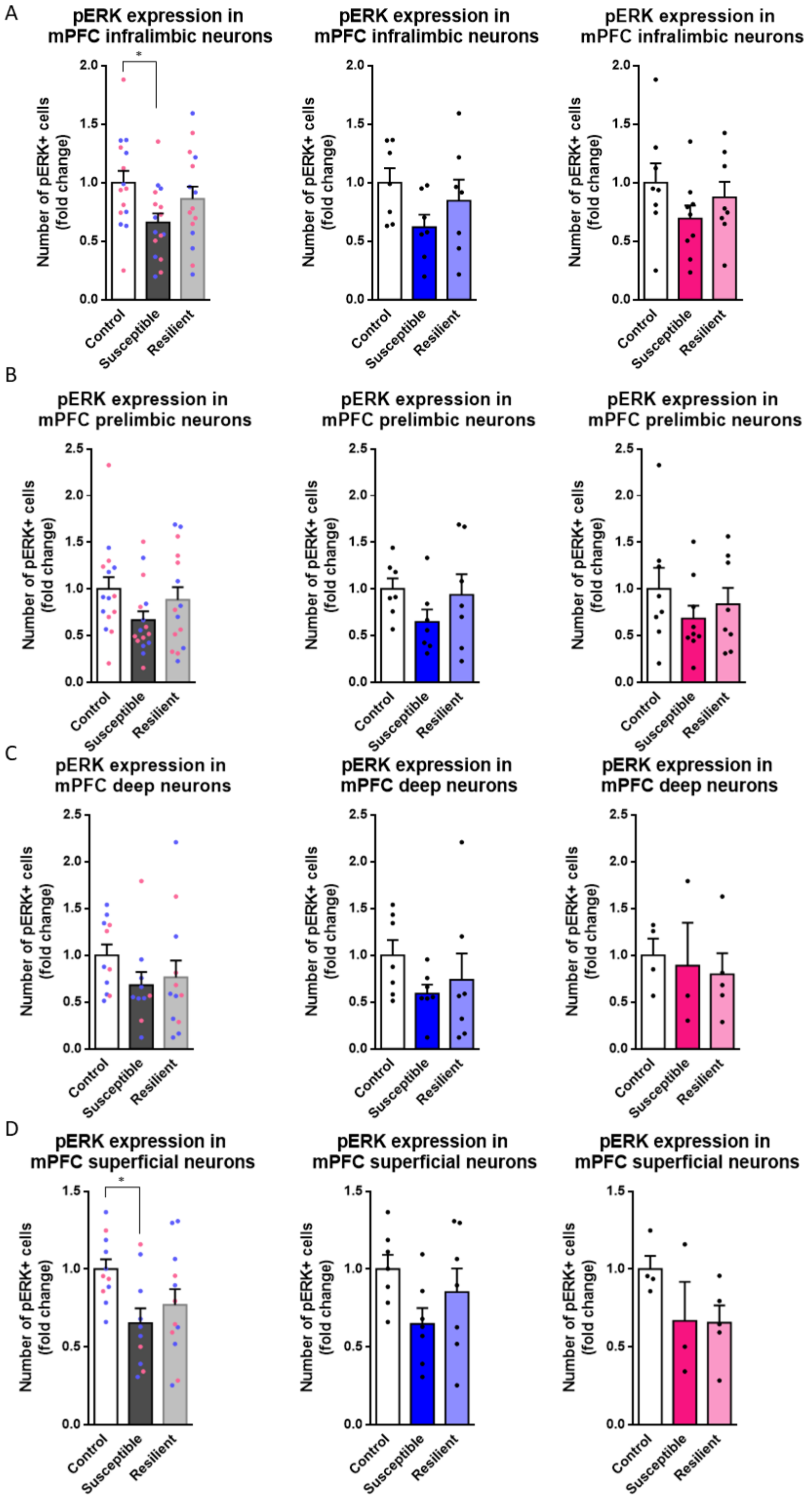

A
pERK expression in mPFC infralimbic neurons
pERK expression in mPFC infralimbic neurons
pERK expression in mPFC infralimbic neurons
B
pERK expression in mPFC prelimbic neurons
pERK expression in mPFC prelimbic neurons
pERK expression in mPFC prelimbic neurons
C
pERK expression in mPFC deep neurons
pERK expression in mPFC deep neurons
pERK expression in mPFC deep neurons
D
pERK expression in mPFC superficial neurons
pERK expression in mPFC superficial neurons
pERK expression in mPFC superficial neurons
Number of pERK+ cells (fold change)
Control
Susceptible
Resilient
*

**<u>Figure S3. pERK expression in the mPFC of males and females after CSDS.</u>** (**A**) Stereological counts of pERK+ neurons in the infralimbic mPFC (females in pink and males in blue [One-way ANOVA, Males and females combined (left panel: $F_{(2,43)}$=3.258; $p$<0.05, Males (middle panel): $F_{(2,18)}$=1.846; $p$=n.s., Females (right panel): $F_{(2,22)}$=1.301; $p$=n.s., Control (M/F): n=7/8, Susceptible (M/F): n=7/9, Resilient (M/F): n=7/8]). (**B**) Stereological counts of pERK+ neurons in the prelimbic mPFC (females in pink and males in blue [One-way ANOVA, Males and females combined (left panel): $F_{(2,43)}$=2.041; $p$=n.s., Males (middle panel): $F_{(2,18)}$=1.354; $p$=n.s., Females (right panel): $F_{(2,22)}$=0.7774; $p$=n.s., Control (M/F): n=7/8, Susceptible (M/F): n=7/9, Resilient (M/F): n=7/8]). (**C**) Stereological counts of pERK+ neurons in deep layers of the mPFC (layer V and VI; females in pink and males in blue [One-way ANOVA, Males and females combined (left panel): $F_{(2,30)}$=1.154; $p$=n.s., Males (middle panel): $F_{(2,18)}$=1.124; $p$=n.s., Female: $F_{(2,15)}$=0.1541; $p$=n.s., Control (M/F): n=7/4, Susceptible (M/F): n=7/3, Resilient (M/F): n=7/5]). (**D**) Stereological counts of pERK+ neurons in superficial layers of the mPFC (layer II/III and I; females in pink and males in blue [One-way ANOVA, Males and females combined (left panel): $F_{(2,30)}$=3.814; $p$<0.05, Males (middle panel): $F_{(2,18)}$=2.233; $p$=n.s., Females (right panel): $F_{(2,9)}$=1.938; $p$=n.s, Control (M/F): n=7/4, Susceptible (M/F): n=7/3, Resilient (M/F): n=7/5]).
